# Dysregulation of Prefrontal Oligodendrocyte Lineage Cells Across Mouse Models of Adversity and Human Major Depressive Disorder

**DOI:** 10.1101/2023.03.09.531989

**Authors:** Sumeet Sharma, Wenjing Ma, Kerry J. Ressler, Thea Anderson, Dan. C. Li, Peng Jin, Shannon L. Gourley, Zhaohui Qin

## Abstract

Animal models of adversity have yielded few molecular mechanisms that translate to human stress-related diseases like major depressive disorder (MDD). We congruently analyze publicly available bulk-tissue transcriptomic data from prefrontal cortex (PFC) in multiple mouse models of adversity and in MDD. We apply strategies, to quantify cell-type specific enrichment from bulk-tissue transcriptomics, utilizing reference single cell RNA sequencing datasets. These analyses reveal conserved patterns of oligodendrocyte (OL) dysregulation across animal experiments, including susceptibility to social defeat, acute cocaine withdrawal, chronic unpredictable stress, early life stress, and adolescent social isolation. Using unbiased methodologies, we further identify a dysregulation of layer 6 neurons that associate with deficits in goal-directed behavior after social isolation. Human post-mortem brains with MDD show similar OL transcriptome changes in Brodmann Areas 8/9 in both male and female patients. This work assesses cell type involvement in an unbiased manner from differential expression analyses across animal models of adversity and human MDD and finds a common signature of OL dysfunction in the frontal cortex.

## Introduction

Psychosocial adversity contributes to multiple neuropsychiatric conditions, including depression, anxiety, addiction, psychosis, and trauma-related disorders^1^. Animal models of stress can reproduce aspects of physiology and partial phenotypes of human stress-related disorders^2^, but they have yielded few molecular pathways that are shared in human disease. Transcriptome studies in animal models implicate diverse aspects of brain function, including the hypothalamic-pituitary axis, neurotransmitters, nuclear receptors, transcriptional regulators, lipid regulation, and others^3–8^. A comparative analysis between chronic variable stress in the mouse and human major depressive disorder (MDD) found little overlap in differentially expressed genes (DEGs) or gene pathways in the prefrontal cortex (PFC). Furthermore, transcriptomes from males and females with MDD had little in common^9^. Identifying conserved molecular features across animal models and human disease remains a major challenge.

Numerous animal models and human disease states have been characterized with unbiased transcriptome sequencing, but few methods exist to find common features across experiments. Bulk tissue transcriptome sequencing has produced an abundance of deeply sequenced data, with large sample sizes that effectively capture individual variation in stress resilience and susceptibility^10–13^. However, individual gene comparisons are limited by sample size and the number of significant DEGs in each experiment, as well as technical and biological variation. Pathway and co-expression analyses of bulk-tissue transcriptomes can identify larger functional units, but these approaches lose cell-type specificity. And while single-cell RNA sequencing (RNA-seq) can pinpoint cell-specific dynamics and unique marker genes, it is often limited by sample size and may not accurately capture changes in cell proportion^1^^4^.

To address some of these gaps in translation, we have created a novel methodology, LRcell, which leverages cell-type specific marker genes derived from single-cell datasets to identify cell-type changes in bulk-tissue RNA-seq experiments. LRcell uses logistic regression to calculate the enrichment of marker genes in summary statistics of bulk-tissue RNA-seq experiments, to produce a set of false discovery rate (FDR) adjusted p-values for each cell type based on differential expression of marker genes. These values can compare relative transcriptional changes by cell type across experiments.

In this work we analyze transcriptomic datasets from animal models of behavioral adversity and human MDD. We focus on the PFC because psychosocial stress has been shown to impair the structure and function of this brain region^15^, and because it is widely transcriptionally characterized. We find signatures of oligodendrocyte (OL) dysregulation across animal models of adversity and human MDD, in both males and females.

## Materials and Methods

### LRcell

LRcell uses logistic regression to assess whether cell-type specific marker genes are differentially expressed in experimental conditions. This requires a list of marker genes for each cell type, identified as previously described^16^, and a list of DEGs ranked by statistical significance. LRcell runs a logistic regression as

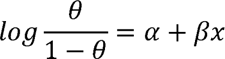

and

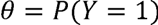

in which *Y* is an indicator variable *Y* = 1 indicates a marker gene and *Y* = 0 indicates otherwise. *θ* thus represents the probability (*P*) that the gene is a marker gene. We use -log(*p* – value) as the explanatory variable *x*. The slope parameter *β* corresponds to the log odds of being one of the marker genes of the cell type.

The contribution of each cell type to overall transcriptional changes can be assessed with the Wald test, using the null hypothesis *β* = 0 or the alternative *β* ≠ 0. LRcell independently analyzes multiple cell types to see which are significantly involved in overall transcriptional changes and produces log-transformed FDR values for each cell type to compare their transcriptional enrichment.

### LRcell validation in matched single-cell and bulk-tissue data

We downloaded pancreatic islet single-cell and bulk-tissue transcriptomes from 4 T2D patients and 3 controls (Table 1). Single-cell counts were pooled across samples and DESeq2 was used to calculate the top 100 marker genes for each cell type following the original cell annotations^17^. Cells with very low counts including MHC class 2 cells (n=5), mast cells (n=7), and endothelial cells (n=16) were excluded. We extracted gene count matrices from bulk tissue data to calculate DEGs between T2D and controls and analyzed these using cell-type specific marker genes with LRcell. We compared LRcell results with cell proportion changes estimated by the MuSiC algorithm^18^, using raw gene counts from bulk-tissue RNA-seq rather than single-cell data as bulk-tissue deconvolution may more accurately represent cell proportions^14^. We ran linear regression as:

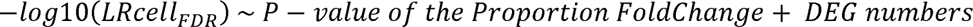

Cell-type specific differential expression of single-cell RNA-seq data was executed using a pseudobulk approach. For each major cell-type identified, single-cell derived counts were aggregated across all cells in that cluster. Then normalization, dispersion estimation, and downstream differential expression analyses were executed using DESeq2^19^, as previously described^20, 21^.

### Cell type proportion estimation

Cell type proportion was estimated by the MuSiC algorithm using default parameters^18^. Cell-specific markers were generated from mouse “Frontal Cortex” single cell data^22^ (Table 1). Multiple replicates were pooled with the *ExpressionSet* object, and read counts of overlapping genes in the bulk datasets were aggregated into another *ExpressionSet* object. The *music_prop()* function calculated cell proportions using *Est.prop.weighted*.

### GO term enrichment analysis

We used the Enrichr program with GO terms in the Molecular Function and Biological Process annotation categories^23, 24^ to identify gene sets shared across stress transcriptomes and within early life isolation data, at FDR<0.1.

### OL lineage trajectory mapping

We classified OL and OPC marker genes as progenitor or mature using the CellMarker database^25^. To chart OL maturation stages from gene expression dynamics, we used Monocle3 to perform UMAP dimension reduction^26^ and create a pseudotime trajectory. Pre-annotated OL and OPC data were extracted and pre-processed by the *preprocess_cds()* function with 100 dimensions, then reduced with the *reduce_dimension()* function of UMAP. We removed one cluster that did not strongly express any known OL or OPC marker genes^25^. The *cluster_cells()* function partitioned the remaining cell populations, and the *learn_graph()* function ordered these populations by pseudotime using gene overlap within and between cell populations, starting with COPs.

### Mouse cell-type specific marker genes

Annotated single-cell data from mouse PFC was downloaded from DropViz, which has synthesized single-cell sequencing data from nine mouse brain regions into 565 cell clusters^22^ (Table 1). We used cluster annotations for the PFC derived from a two-stage Independent Component Analysis (ICA), as previously described^22^. Inactive genes (expressed in less than 10 cells) and inactive cells (expressing less than 10 genes) were excluded from single-cell analyses.

### Mouse models of adversity

Stress-related behavioral paradigms were found by searching GEO^27^ for the terms ‘stress’, ‘mouse’, and ‘prefrontal cortex’. We used datasets showing behavioral changes from adversity that had ≥20 million reads per sample and n>3 per condition. Datasets that met these criteria represented chronic social defeat^4^, early-life stress (maternal separation with limited bedding)^7^, cocaine withdrawal^3^, and chronic variable stress^5^. Raw fastq files were downloaded from SRAdb (Table 1).

We standardized the alignment and annotations of raw reads using STAR alignment software^28^ (options: --quantMode GeneCounts --outSAMstrandField intronMotif -- outFilterIntronMotifs RemoveNoncanonical). Un-normalized counts were input intoDESeq2^29^ for differential expression. We maintained group designations and covariates from the original datasets when applicable.

### Mouse models of adversity

Chronic social defeat^4^:

Early-life stress (maternal separation with limited bedding)^7^:

cocaine withdrawal^3^:

chronic variable stress^5^:

### Social isolation and reward devaluation

#### Subjects

Female C57BL/6 mice from Jackson Labs stock (Bar Harbor, ME) were bred in-house. Socially isolated mice were weaned at postnatal day (P) 21-22 and housed with siblings until single-housing from P31-P60 as previously described^30^. Then, mice were reintegrated into social housing, with each cage consisting of ½ mice that were previously isolated. All mice were provided food and water *ad libitum* except during instrumental conditioning, when they were food-restricted to ∼90-93% of their body weight. Mice were maintained on a 12:12-hour light cycle (0700 on). Procedures were in accordance with the Emory University IACUC.

#### Response training and devaluation

Mice were trained to nose-poke for a reward as previously described^30^, first on a fixed ratio 1 (FR1) schedule of 30 pellets at each aperture and then on a random interval (RI) 30-sec schedule. To test general behavioral flexibility, reinforcement shifted to a previously unrewarded center aperture on an FR1 schedule, so that mice had to inhibit their previous responses and produce a new response. Sessions were 25 min long. To test reward valuation, mice were given *ad libitum* access to the reward pellets for 1 hour to reduce their value. Mice in the original reward condition were pre-fed with chow to control for satiety. Reward devaluation tests were 25 min in extinction conditions, and the order of conditions was randomized.

As previously described^31^, mice were trained on an RI-30-sec reinforcement schedule for 4 additional sessions and then on an RI-60-sec schedule for 2 sessions to induce habitual responses. The devaluation procedure was repeated as described above. Our goal was to assess whether mice with a history of isolation were predisposed towards habit-based responses.

### Tissue processing and RNA-sequencing

#### Orbitofrontal cortex tissue collection

After the second devaluation test, mice were briefly anaesthetized with isoflurane and decapitated. Methods are as previously described^32^, but briefly: brains were rapidly extracted and frozen at –80°C. An investigator blind to condition sectioned fresh-frozen brains on the Microm HM 450 Sliding Microtome with Tissue-Tek O.C.T. compound with the tissue kept at −23.0°C. Tissue punches were taken from the ventrolateral orbitofrontal subregion of the PFC using a 1.0 mm punch biopsy tool. Punches were stored at – 80°C for RNA-sequencing.

#### RNA Isolation and Sequencing

RNA-seq library construction was performed using the Illumina mRNA sample prep kit (cat. no. RS-100-0801). Briefly, we purified poly-A-containing mRNA using poly-T bound magnetic beads. mRNA was fragmented using divalent cations under high temperature. Cleaved RNA fragments were copied into first-strand and then second-strand cDNA synthesis using reverse transcriptase and random primers, and DNA Polymerase I and RNase H, respectively. cDNA fragments were end-repaired by addition of a single ‘A’ base and then ligation of sequencing adapters. The subsequent products were gel purified and analyzed using a bioanalyzer to verify the size and concentration before sequencing on the Illumina HiSeq 2500.

#### Statistics for behavioral and immunostaining studies

Response rates, food consumed, response ratios, and immunofluorescence were compared by ANOVA, with repeated measures as appropriate, or t-tests. Post-hoc comparisons were applied following interaction effects, and p≤0.05 was considered significant.

## Results

### Logistic regression of cell-type specific marker genes correlates with changes in cell proportion

Before analyzing brain transcriptomes, we identified a pancreatic dataset for which bulk-tissue and single-cell RNA sequencing data from the exact same samples are available to identify the transcriptional characteristics represented by LRcell. While many datasets contain single cell and bulk-tissue RNA-seq analyses, few studies specify that samples for bulk RNA-seq were obtained from the same aliquots used for single cell analysis. We compared single-cell and bulk-tissue RNA-seq data of pancreatic islets from patients with type 2 diabetes (T2D) or matched controls^17^. We reasoned that enrichment of cell-type specific marker genes in bulk DEGs may indicate expression changes within cell types or proportion changes between cell types.

LRcell analysis of bulk-pancreas RNA-seq between T2D and controls highlighted enrichment of ductal, pancreatic stellate (PSC), and beta cells (Figure 1A). We compared these results to estimates of cell proportion changes^18^, which showed alterations in ductal and PSC cells that approached significance (ductal 1.897-fold increase, p=0.06; PSC 1.557-fold increase, p=0.056; Figure 1B). We next used a pseudobulk approach^20, 21^ to identify DEGs in single-cell data (Figure 1C). The LRcell signal was significantly correlated with modeled interactions between cell proportion changes and DEGs (methods), with significant *post hoc* correlation between cell proportion changes and the LRcell-derived p-value (p=0.0013; Figure 1D).

**Figure 1.**
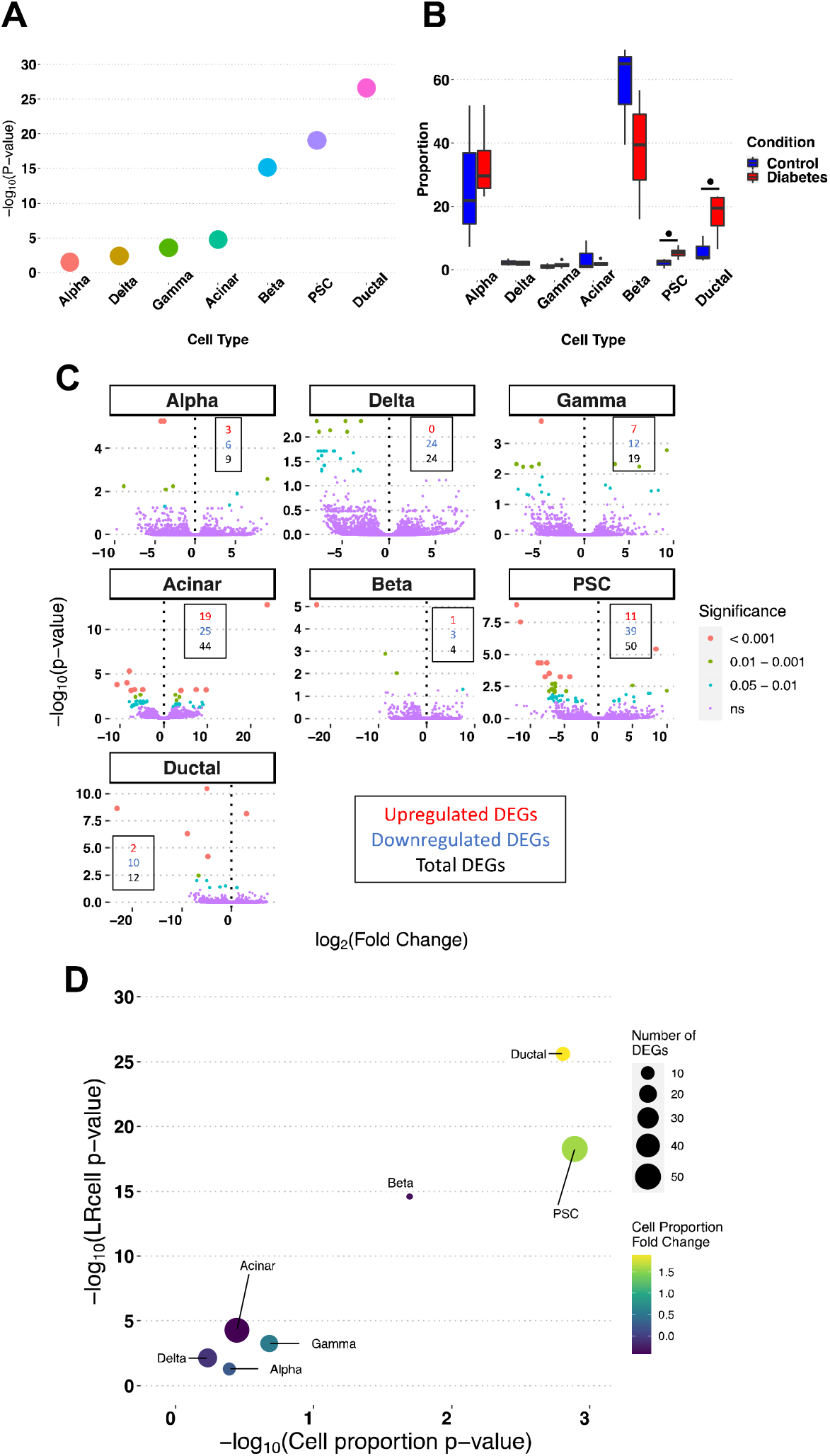
Comparative analyses of matched single-cell and bulk-tissue RNA-seq from T2D and control pancreatic tissue. **(A)** LRcell analysis of pancreatic DEGs between T2D and matched controls. **(B)** Sample-level pancreatic cell type proportions in between T2D and controls (^•^P <0.1). **(C)** Statistical significance (y-axis) and fold change (x-axis) of DEGs in each cell type. Color and size of gene points indicate their statistical significance. Boxes list the number of significant (FDR<0.05) upregulated (red), downregulated (blue) and total (black) DEGs for each cell type. **(D)** LRcell correlation with cell proportion changes and DEGs within cell types: y-axis shows LRcell p-values, x-axis shows p-values of cell proportion changes, dot color scale shows cell proportion fold-change, and dot size shows number of DEGs in each cell type. Overall model statistics: F = 31.77, R^2^ = 0.91, *P* = 0.0035; significant *post hoc* correlation with changes in cell proportion, t-value = 7.971, *P* = 0.0013.

### LRcell implicates prefrontal oligodendrocyte lineage cells and myelination pathways across adversity models

Next we used LRcell to compare prefrontal cortical cell-type specific changes in mouse models of adversity including chronic social defeat^4^, early-life stress (maternal separation with limited bedding)^7^, cocaine withdrawal^3^, and chronic variable stress^5^ (Table S1). We identified cell-type specific marker genes in PFC from publicly available single-cell RNA-seq data^22^ (Figure 2A). DEGs varied across experiments, but LRcell analysis of all adversity models showed consistent marker gene enrichment of mature oligodendrocytes (OLs) and oligodendrocyte precursor cell (OPCs) (Figure 2B). We further corroborated these results with an independent single cell reference dataset^33^, observing similar enrichment of mature OL-lineage cells (Figure S1). Social defeat led to enrichment of mature OL and OPC markers in behaviorally susceptible animals but not in resilient animals (Figure S2A), with minimal signal from other cell types. Transcriptomes from acute cocaine withdrawal, long-term cocaine withdrawal, and cocaine administration showed enrichment of OL and OPC marker genes only in acute withdrawal, the most aversive condition (Figures 2B, S2B, S2C). LRcell analysis of acute withdrawal also highlighted vascular endothelia, vascular smooth muscle cells, and astrocytes, which are all sensitive to the sympathomimetic effects of cocaine or previously implicated in withdrawal^34^ (Figure 2B). Chronic unpredictable stress induced significant transcriptional differences in OL marker genes but also showed broad enrichment of interneurons, OPCs, claustrum, layers 6 and 2/3 excitatory neurons, and astrocytes. Subsets of neurons from this condition exhibited greater marker gene dysregulation than other cell types: a subset of layer 2/3 neurons showed greater transcriptional differences than any other cell type including OLs while the remaining layer 2/3 neurons were only modestly affected (Figure 2B). LRcell analysis of early life stress showed marker gene enrichment from OLs and OPCs, but interestingly, when mice with a history of early life stress experienced social defeat in adulthood they showed enrichment of brain fibroblast-like cells without a significant OL signal (Figures 2B, S2D).

**Figure 2.**
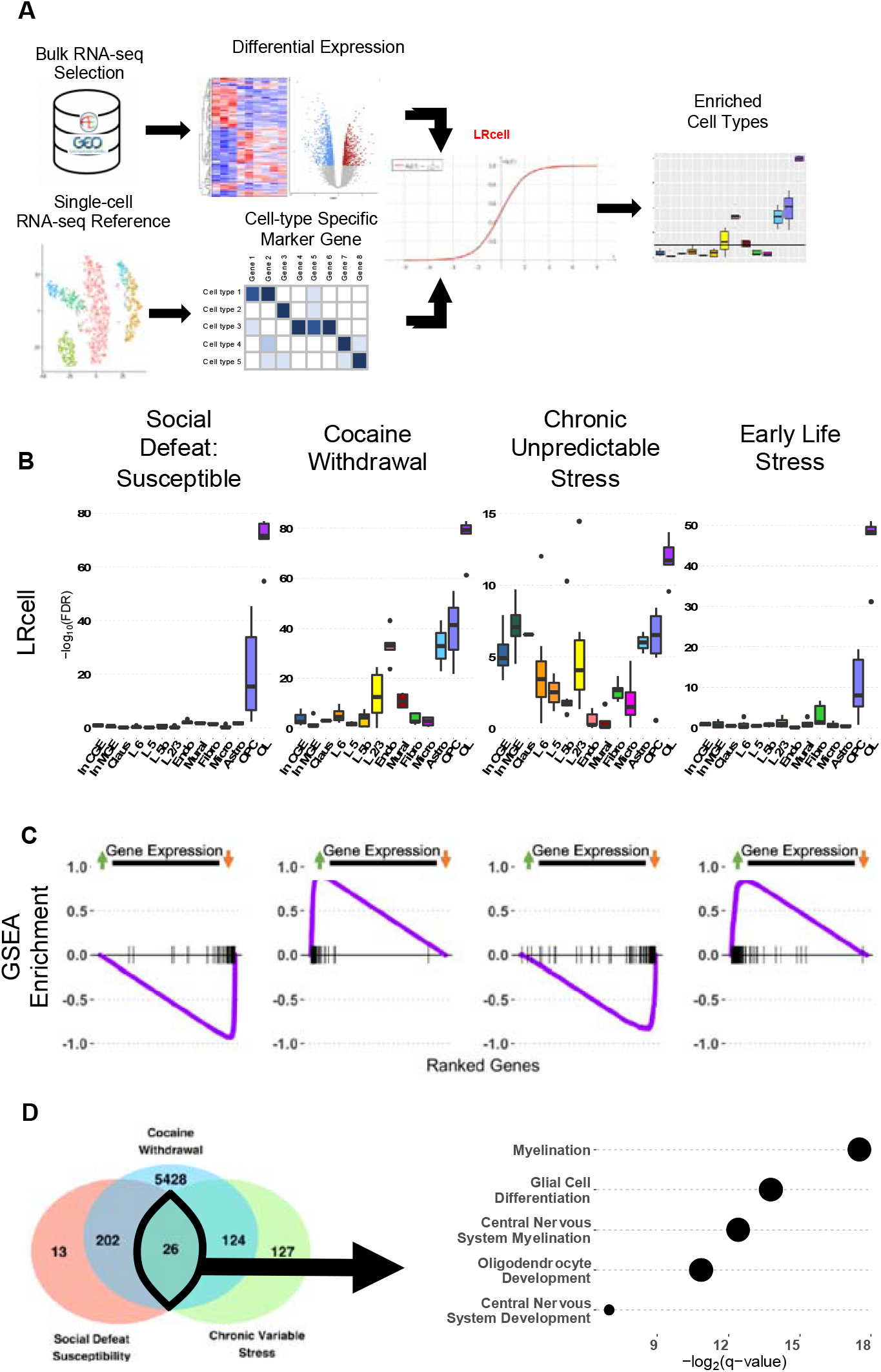
Convergent analysis of mouse models of adversity. **(A)** Schematic of analytical strategy. **(B)** LRcell analyses of four adversity models. Box plots depict FDR-adjusted p-values for all single-cell RNA-seq defined cell types within each cluster. **(C)** Enrichment of OL marker genes ranked on a continuum by -log_10_(p-value) x sign(log_2_(fold-change)): significantly upregulated genes to the left, and significantly downregulated genes to the right. **D) Left**: Specific and overlapping DEGs in each experiment (FDR-adjusted p-value < 0.1). **Right**: Pathway analysis of DEGs significant in all three datasets. Dot size is proportional to p-value. ***Abbreviations:* In CGE; In MGE**: interneurons from caudal or medial ganglion eminence; **Claus**: claustrum; **L6**, **L5**, **L5b**, **L2/3**: cortical layers 6, 5, 5b, 2 and 3; **Endo**: endothelial cells; **Mural**: mural cells; **Fibro**: fibroblast-like cells; **Micro**: microglia; **Astro**: astrocytes; **OPC**: oligodendrocyte precursor cells; **OL**: mature oligodendrocytes.

We estimated OL-lineage proportion changes^18^ and assessed directional changes in marker gene expression in these models of behavioral adversity. OL and OPC proportions were significantly increased in in cocaine withdrawal (Figure S3). No other conditions showed significant OL-lineage proportion changes at p<0.05, but the cocaine withdrawal dataset had by far the largest sample size (n ∼ 50). Our power calculations indicated that to detect proportion differences at a significance level of 5% and power of 80%, average sample sizes of 17 are required for OL and 30 for OPC populations. Enrichment plots of the mature OL lineage marker genes showed that transcriptional deviations were either consistently upregulated or downregulated (Figure 2C). In the susceptibility to social defeat and chronic unpredictable stress paradigms, mature OL markers were downregulated compared to controls, while in the early life stress and cocaine withdrawal experiments, mature OL markers were upregulated.

Lastly, we used Enrichr with Gene Ontology (GO) annotations^23, 24^ to identify biological pathways shared across stress paradigms. The early life stress dataset had no DEGs that met the criteria of FDR<0.1 so it was excluded. Twenty-six DEGs were shared across social defeat, cocaine withdrawal, and chronic variable stress (Figure 2D: Left). These DEGs were enriched in myelination, glial development, and CNS development pathways (Figure 2D: Right).

### Early life isolation disrupts goal-directed behavior, oligodendrocytes, and layer 6 neurons

Our initial findings suggest that genes specific to OL-lineage cells are consistently and disproportionally affected by stress and adversity. These observations led us to the testable hypotheses that other adverse experiences would similarly affect OL gene expression, that expression changes might correlate with behavioral changes from adversity, and that these changes might be consistent across both sexes. To test these possibilities, we used a model of social isolation that we have previously described^30^. In short, female mice were isolated from postnatal day (P)31 in adolescence until P60 in young adulthood, then tested for long-term neurobehavioral effects (Figures 3A and 3B).

**Figure 3.**
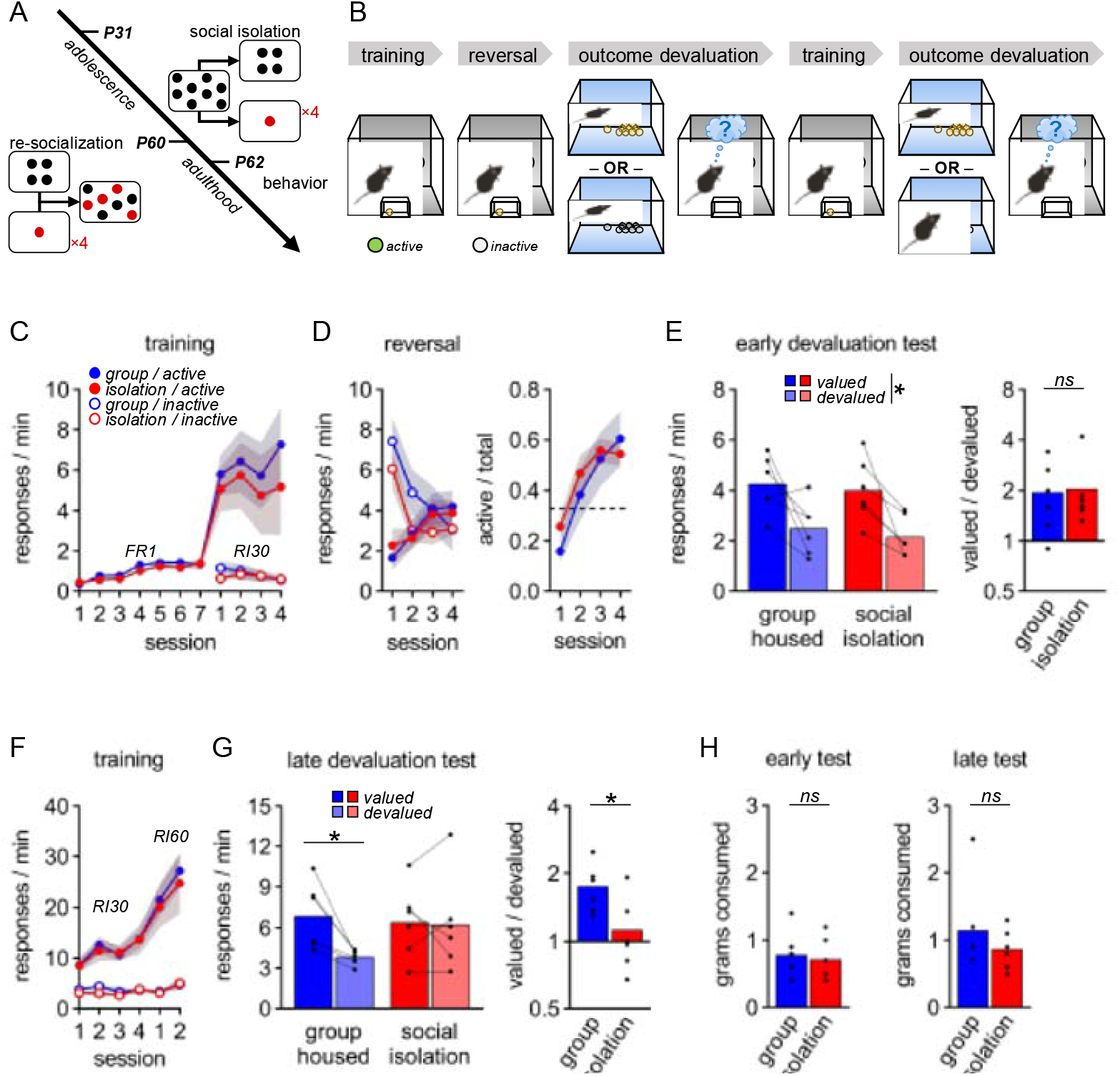
Social isolation impairs reward valuation in goal-directed behavior. **(A)** Experimental timeline: social isolation from P31-P60 and behavior testing beginning at P62. **(B)** Schematic of behavioral tasks. **(C)** Initial response training. Training progressed from an FR1 schedule of reinforcement (session: F_6,60_ = 19.49, p < 0.01; no effect of isolation or session × isolation: F < 1) to an RI30 schedule (session × aperture: F_3,30_ = 1.29, p = 0.30; main effect of aperture F_1,10_ = 26.74, p < 0.01; no effect of isolation or session × aperture × isolation: F < 1). **(D)** Instrumental reversal. **Left:** Responses to new reward locations indicate that both groups could modify response patterns (session × aperture: F_3,30_ = 16.35, p < 0.01; no effect of isolation or session × aperture × isolation: F < 1). **Right:** Reversal rates (active / total nose-pokes) across sessions exceeded chance (0.33) in both groups (session: F_3,30_ = 38.79, p < 0.01; session × isolation: F_3,30_ = 1.86, p=0.16; no main effects: F=1). **(E)** All mice inhibited responses to a devalued reward (**left:** raw response rates, main effect of outcome condition: F_1,10_=18.34, p < 0.01; no effect of isolation or outcome × isolation interaction: F < 1). **Right:** Preference ratios (valued / devalued condition) did not differ between groups (t_10_ < 1). **(F)** Additional training with RI reinforcement schedule (session × aperture: F_5,50_=20.41, p < 0.01; main effect of aperture: F_1,10_=62.50, p < 0.01; no effect of isolation or session × aperture × isolation: F < 1). **(G)** Reward devaluation test after RI training shows that socially isolated mice fail to modify responses based on outcome value. **Left:** Raw response rates (main effect of outcome: F_1,10_=6.54, p < 0.05; outcome × isolation: F_1,10_=5.24, p < 0.05; no effect of isolation: F < 1). **Right:** Control mice responded more in valued condition whereas previously isolated mice did not change their response rate based on outcome value (between groups t_10_=2.43, p < 0.05). **(H)** All groups consumed the same amount of food before testing.

Isolation did not affect general reinforcement learning (Figure 3C) or reversal learning (Figure 3D). Group-housed and socially isolated mice also similarly adjusted their response rate when the reward was devalued through *ad libitum* access (Figure 3E), demonstrating similar assessment of value. However, when isolated animals were switched to random interval (RI) reinforcement, which induces more habitual responses over time (Figure 3F), they became insensitive to reward devaluation while the group-housed mice were still able to adjust their behavior to reward value (Figure 3G). Group differences did not reflect differences in food consumption before testing (Figure 3H).

LRcell analysis of transcriptomes from the PFC of socially isolated and group-housed animals showed significant differences in OLs and OPCs, and in layers 6 and 2/3 excitatory neurons (Figure 4A Left). Mature OL marker genes were upregulated (Figure 4A Right) and GO term enrichment analysis highlighted cellular development and differentiation, myelination, and axon development (Figure 4B), like other adversity models (Figure 2D). OL cell proportion in socially isolated animals trended towards an increase compared to controls (p=0.06; Figure 4C).

**Figure 4.**
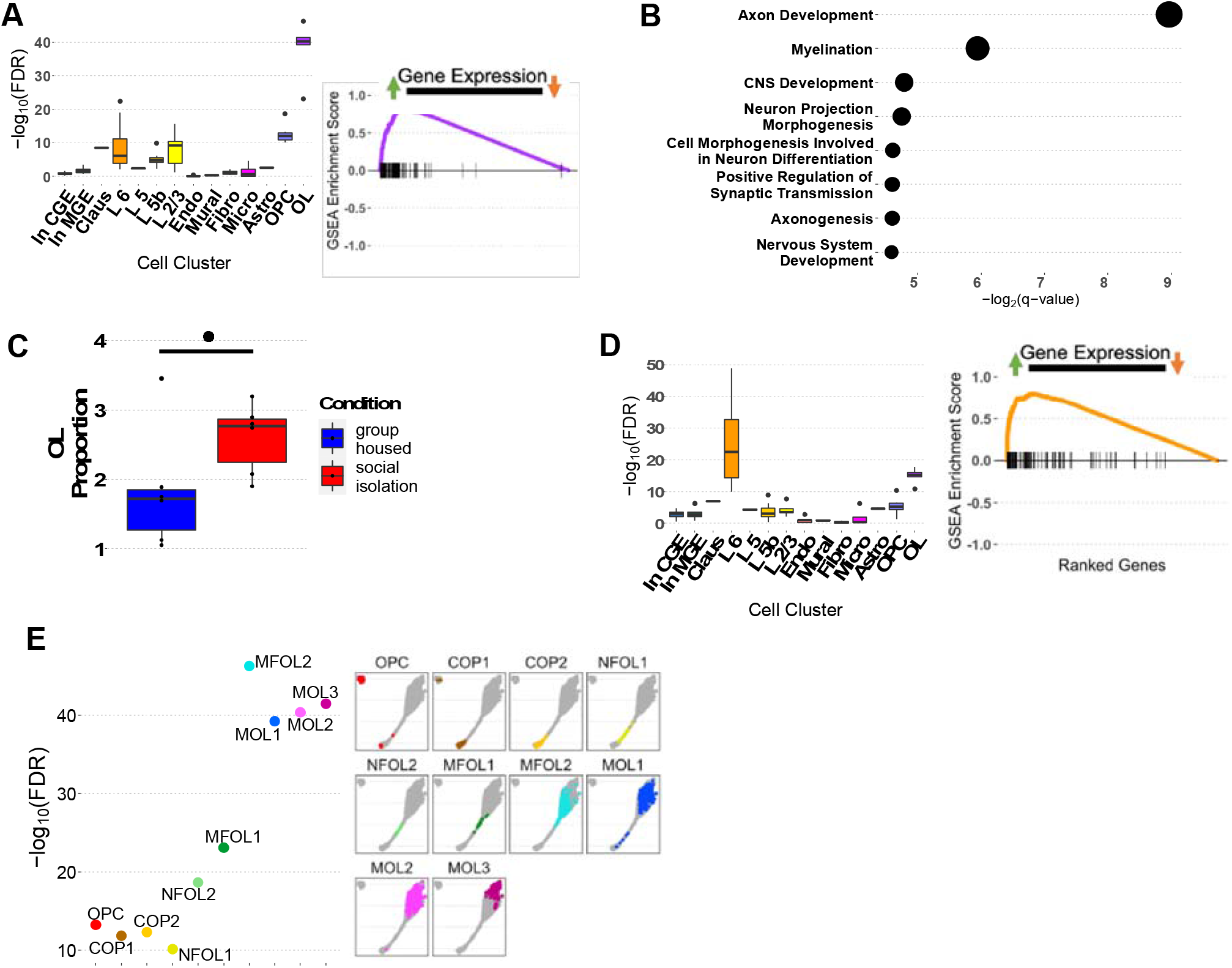
Transcriptional enrichment of OL-lineage cells in adolescent social isolation. **(A) Left:** LRcell analyses of DEGs comparing socially isolated and group housed animals (n=6). **Right:** OL marker gene enrichment between socially isolated and group-housed animals. **(B)** Pathway analysis of DEGs from social isolation; dot size indicates p-value. **(C)** Comparison of oligodendrocyte cell % in group-housed and isolated animals (Wilcox rank sum test, p-value = 0.06, W = 6; ^•^ P <0.1). **(D)** Comparison of highest and lowest responding tertiles after reward devaluation (n=4). **Left:** LRcell analysis of DEGs, and **Right:** Layer 6 excitatory neuron marker gene enrichment. **(E) Left:** Expanded LRcell analyses of OPC and OL cell types, ordered by developmental stage. **Right:** UMAP representation of OL cell types, with shaded region representing entire OL and OPC population and colored points corresponding to cell type. ***Abbreviations:* In CGE; In MGE**: interneurons from caudal or medial ganglion eminence; **Claus**: claustrum; **L6**, **L5**, **L5b**, **L2/3**: cortical layers 6, 5, 5b, 2 and 3; **Endo**: endothelial cells; **Mural**: mural cells; **Fibro**: fibroblast-like cells; **Micro**: microglia; **Astro**: astrocytes; **OPC**: oligodendrocyte precursor cells; **COP**: committed oligodendrocyte precursors; **NFOL**: newly formed oligodendrocytes; **MFOL**: myelin forming oligodendrocytes, **MOL** or **OL**: mature oligodendrocytes.

Early social isolation biases animals toward habitual behavior, but individual animals still show a range of behavioral sensitivity to reward devaluation. To investigate possible associations between behavioral phenotype and cell type enrichment, we ranked animals by their response ratio (responses in valued condition ÷ total responses). A higher response ratio indicates more goal-directed behavior and greater sensitivity to reward value. We compared transcriptomes between the lowest and highest-ranked tertiles (n=4). LRcell analysis of DEGs from this comparison highlighted layer 6 excitatory neurons (Figure 4D Left), and we saw an upregulation of layer 6 marker genes in animals with the most impaired goal-oriented behavior (Figure 4D Right). OLs were also affected, but less than in comparison between all isolated and control animals.

### Behavioral adversity affects maturing oligodendrocytes

OPCs mature through distinct stages as they differentiate, find a niche, and initiate and complete myelination^34^, so we further annotated OL-lineage cells to examine transcriptional changes during maturation. We labeled genes within clusters as progenitor or mature using the CellMarker database^25^, then putatively ordered cell subtypes by maturation stage based on label proportions (Figure S4A). In a separate analysis, we used UMAP dimension reduction to identify one cluster of pure OPCs and a separate cluster of heterogeneous OL-lineage cells in different stages of maturation. Trajectory analysis of these OL-lineage subtypes confirmed our initial staging (Figure S4B, S4C)^22, 26^. We characterized these subclusters as OPCs (oligodendrocyte precursor cells), COPs (committed oligodendrocyte precursors), NFOLs (newly formed oligodendrocytes), MFOLs (myelin forming oligodendrocytes), and MOLs (mature oligodendrocytes) according to well-characterized gene expression at each stage^35^ (Figure S5).

We used LRcell with these annotations to compare transcriptional changes in OL maturation stages after social isolation. The most mature OL cell-types (MOL1-3 and MFOL2) were enriched among ranked bulk-tissue DEGs, intermediate populations (NFOL2 and MFOL1) showed intermediate enrichment, and precursor populations (OPC, COP, NFOL1) were the least enriched of OL lineage cells (Figures 4E). Other adversity paradigms showed similar differential enrichment of OL-lineage cells that increased with maturation stage (Figure S6). To ensure that this pattern did not simply reflect similarities in marker gene sets, we examined marker gene correlation among all cell types in the mouse PFC. Marker genes of mature OL populations were highly correlated with each other but distinct from intermediate and earlier OL maturation stages (Figure S7A).

### Mature oligodendrocytes are dysregulated in major depressive disorder

Lastly, we examined cell-type specific signatures in human bulk-tissue transcriptomes from patients with MDD. We used a prefrontal cortical single nucleus RNA-seq study of MDD^36^ to identify cell-type specific marker genes in Brodmann Areas 8/9 (Figure 5A). Marker gene identities were minimally impacted by MDD status (Figure S7B), so we derived marker genes for LRcell using control samples. Some marker genes in Brodmann Areas 8/9 were conserved with those in mouse PFC, with the most overlap in OLs, other glia, and interneurons and the least overlap in cortical excitatory neurons (Figure 5B). Maturation stages inferred by the overlap between mouse and human OL marker genes were consistent with trajectory analysis^36^.

**Figure 5.**
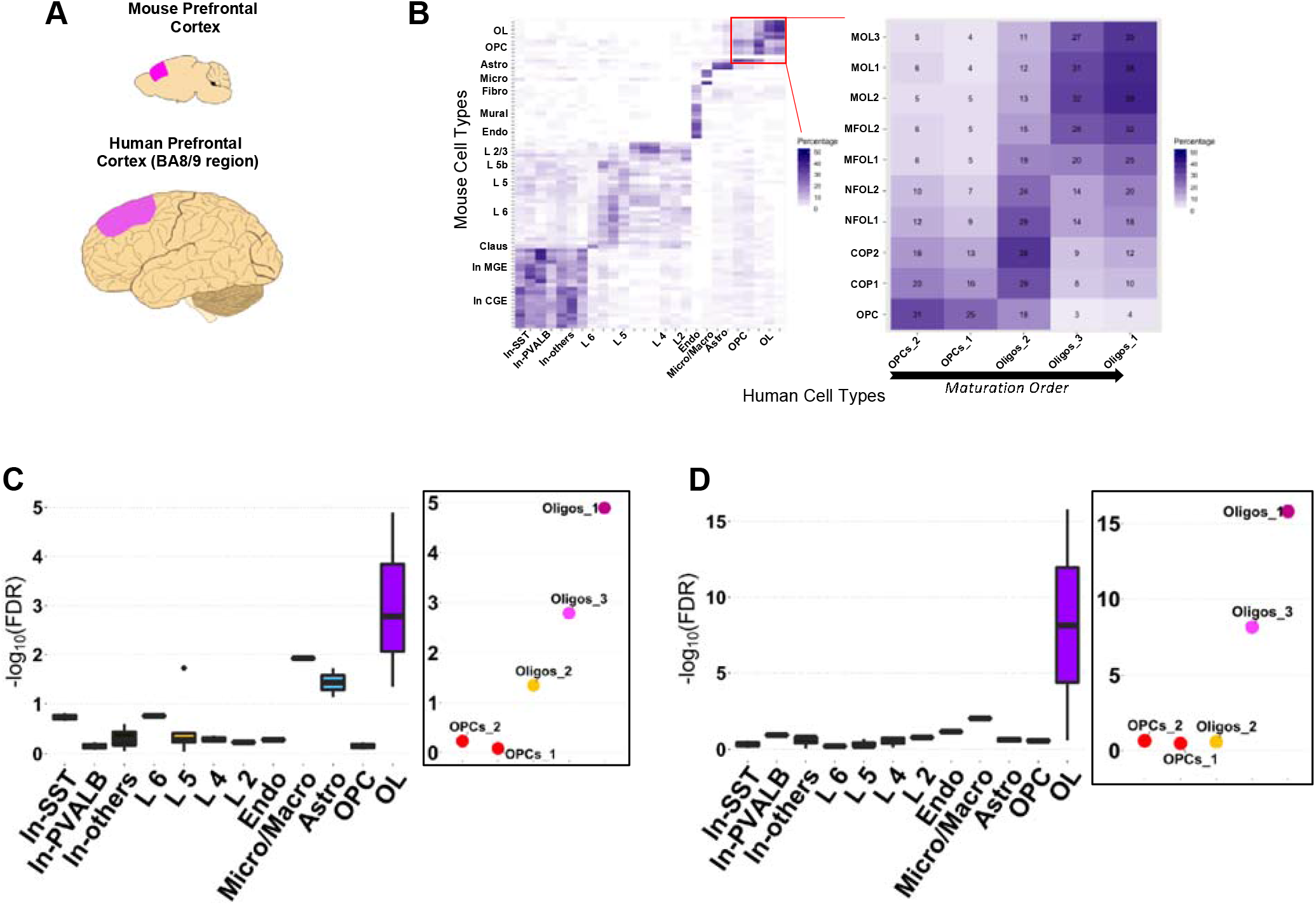
LRCell analyses of male and female post-mortem samples from human major depressive disorder. **(A)** Brain regions investigated in mouse adversity paradigms and human MDD. **(B) Left**: Overlap between mouse prefrontal cortical cell-type markers and human BA8/9 cell markers, with major cell type clusters labeled. Mouse genes annotations (mm10) were lifted over to human (hg38). **Right**: Expansion of the OL and OPC cell populations. **(C) Left:** LRcell analysis of DEGs from male MDD compared to control in BA8/9 using single-cell references from BA9 region. **Right:** Expansion of LRcell analyses of OPC and OL cell types ordered by lineage trajectory. OL cluster labels (Oligos_2, Oligos_3, Oligos_1) are categorical. **(D)** Corresponding LRcell analyses in female MDD. ***Abbreviations:* In-SST**, **In-PVALB, In-others**: interneurons expressing somatostatin, parvalbumin, or neither; **L6**, **L5**, **L5b**, **L4, L2/3**: cortical layers 6, 5, 5b, 4, 2 and 3; **Endo**: endothelial cells; **Micro/Macro**: microglia or macrophage lineage cells; **Astro**: astrocytes; **OPC**: oligodendrocyte precursor cells; **OL**: mature oligodendrocytes.

LRcell analysis of bulk transcriptomes from human MDD^9^ highlighted mature OL differences in both males and females. Using the original publication’s covariates, we separately calculated DEGs for males and females, adjusting for age, RNA integrity number (RIN), alcohol use, and medication status^9^. Mature OL populations were most enriched in both male and female MDD (Figures 5C Left, 5D Left). OPC and OL maturation stages showed a similar pattern to those of animal models, with LRcell enrichment highest in mature OL populations (Figures 5C Right, 5D Right).

## Discussion

### Future directions for cell-type deconvolution methods

Recent transcriptome deconvolution techniques use cell-specific signatures to estimate cell proportions from bulk-tissue RNA-seq data^14, 18^. LRcell takes an alternative approach by identifying cell type-specific enrichment from differential expression analyses of bulk-tissue RNA-seq. By leveraging differential expression data, LRcell highlights cell types with significant transcriptional alterations between experimental groups.

We recognize several limitations of LRcell and related deconvolution approaches. First, these programs require accurate marker gene lists for the tissues of interest. Although new single-cell technologies and atlases^37, 38^ make these lists increasingly accessible, more work is needed to achieve consensus datasets. Second, the overlap of marker genes across similar cell types can influence the LRcell enrichment calculation. In this work we used all available cell types from the single-cell datasets we selected, which could lead to correlated results but also enables more granular detection. For instance, the differences we observed in intermediate OL maturation stages (NFOL1-2, MFOL1) may have been obscured if cells were simply categorized as OPCs or OLs. These correlations can be mitigated by using tree-based methodologies or adjustable merging of cell types^18^. And thirdly, LRcell cannot detect expression changes in genes common to multiple cell types. Future work may aim to further identify cell-type specific variant genes and incorporate prior knowledge of their dynamics to improve cell-type specific deconvolution of bulk RNA-seq data.

### Multiple types of adversity disrupt OL development

In this work we combine multiple high-quality PFC transcriptomic studies in animal and human studies to identify cell type specific enrichment. While a multitude of previous hypothesis directed work has examined cell-type specific dynamics in stress models, we have developed LRcell to generate an unbiased analysis of all cell types identified by single cell RNA-seq. Our results suggest OL-lineage cell types are the most transcriptionally altered by the experience of psychosocial adversity across animal models and in human MDD.

Animal stress models and human neuropathology have shown changes in OLs and white matter from behavioral adversity^29, 39–41^. Early life stress impairs OL process formation, branching complexity, and neuronal contacts^29^. These morphological defects accompany differences in cell proportion^42^, with depleted OPCs and increased mature OLs^29^, raising the possibility that that early life stress accelerates OL maturation at the expense of thorough myelination. Consistent with this possibility, our results show widespread prefrontal cortical OL dysregulation across animal models of adversity, and that this dysregulation increases as cells mature.

Although OL transcriptional signatures were consistently disrupted in our analyses, the direction of these changes varied by adversity model. Marker genes were upregulated in cocaine withdrawal and early life stress but downregulated in animals susceptible to social defeat and after chronic unpredictable stress (Figure 2B). This suggests that different types of stress, or susceptibility to stress, may interact with OL biology through distinct mechanisms.

Existing studies of early life social isolation have largely focused on myelin dynamics shortly after the isolation period, both in adult and early life isolation models. qPCR-based quantification of candidate oligodendrocyte-specific genes show reduced gene expression during or at the end of the isolation period^29, 43^. In this work, we observe upregulation of single-cell defined OL-lineage genes and higher numbers of mature OLs in adult animals with histories of adolescent social isolation and early life stress (Figure 2B). This reflects recent work that has shown that accelerated OPC maturation occurs as a consequence of early life stress^44^. Taken together these results suggest that adverse early life social experience results in an initial downregulation of OL-specific genes during or shortly after adolescent social isolation, and an upregulation of these genes in adulthood, likely as a consequence of an accelerated maturation of OPCs.

### Transcriptional changes in cortical layer 6 correlate with impairments in goal-directed behavior

In our analysis, layer 6 excitatory neurons were significantly enriched and upregulated in animals with the most habitual behavior after social isolation compared to those with the most behavioral resilience. Previous hypothesis-driven work has similarly shown that early life social isolation alters deep layer neurons^39^, and preferentially thalamocortical projection neurons to impair behavior^29, 45, 46^. In human studies, stress predisposes individuals toward habitual behaviors that are insensitive to goals^47^, which likely contributes to increased obesity, smoking, and other negative responses associated with stress. Yet humans and rodents alike show a broad range of vulnerability or resilience to stress. Transcriptomes comparing our most behaviorally susceptible and resilient animals differed most significantly in layer 6 neurons, suggesting that these neuronal populations may mediate impairment or resilience in goal-directed behavior after stress.

The relationships between pathophysiological neuronal activity and corresponding OL dysfunction are not clear. Research across human psychiatric diseases shows fewer OLs in deep cortical layers in MDD, bipolar disorder, and schizophrenia^48, 49^. And further ultra-structural analyses of myelination in the mouse show the highest density of myelin in deep layer excitatory neurons^50^, suggesting these may be susceptible to myelination deficits. Future work should further examine how dysregulation of deep layer neuronal circuitry affects OL physiology.

### Myelin and OL-lineage dynamics in human MDD

Stress can dysregulate myelination in animal models and in human disorders like PTSD and MDD^51^. Decades of brain imaging show strong associations between white matter disturbances and depression.^52^ Likewise, in this work we observe mature OL dysregulation across multiple animal adversity models and in men and women with MDD.

Recent single-cell sequencing from MDD found the most cell-type specific DEGs in OPCs rather than mature OLs^36^ and there are several possible interpretations of this difference. LRcell highlights cell proportion driven effects that single cell studies may not recognize^14^, but does not detect differences in broadly expressed genes. The DEGs most strongly expressed in OPCs may be common to other cell types. While OPCs may harbor many cell-type specific alterations in MDD, the net result may be a dysregulated maturation of OLs. Other sources of discrepancy may be cohort effects, given the relatively small sample size of both studies, or technical differences in the analytical pipelines for single-cell and bulk-tissue RNA-seq approaches.

Human oligodendrocyte dynamics are still not well understood. Previous work has suggested that human OLs are stable in adulthood with little active turnover^53^, whereas OPCs and OLs in mice show plasticity, division, and maturation even in discrete learning events^54^. Our work suggests that MDD and animal adversity models converge at OL pathophysiology, but translational researchers should be cautious in extrapolating specific physiological or therapeutic effects from mouse to human.

Our findings suggest that deficits in the OL lineage may mediate long-lasting neurobiological changes from stress, and that these deficits are shared across genders, in animal models of adversity and human MDD. We further present a robust and easy to use new methodology, LRcell, to elucidate cell-type specific signals from bulk-tissue transcriptomic analyses.

## Acknowledgements

The authors thank Elizabeth Hinton and Aylet Allen for important contributions to *in vivo* experiments. This work was supported by NIH MH117103, OD011132

## Author Contributions

Designed research (SS, WM, SG, ZQ), performed research (SS, WM), analyzed data (SS, WM, DL, TA), contributed unpublished reagents/analytic tools (DL, PJ, SG, KR), wrote the paper (SS, WM, TA, KR, SG, ZQ)

---

**Table S1.** Experiments analyzed in the manuscript.

|  | Sex | Sample Size | Species | Data Type | Accession |
| --- | --- | --- | --- | --- | --- |
| Chronic social defeat | Male | n = 4 | Mouse | Bulk tissue | SRP062829 |
| Cocaine self-administration & withdrawal | Male | n = 5-8 | Mouse | Bulk tissue | SRP132477 |
| Early life stress | Male | n = 4-6 | Mouse | Bulk tissue | SRP093288 |
| Chronic unpredictable mild stress | Male | n = 2 | Mouse | Bulk tissue | SRP075361 |
| Single cell atlas | Male | - | Mouse | Single cell | SRP151685 |
| Single cell analysis of PFC | Male | - | Mouse | Single cell | SRP178464 |
| MDD | Male, Female | n = 9-13 | Human | Bulk tissue | SRP115956 |
| Type 2 Diabetes | Male, Female | n = 3-4 | Human | Single cell, bulk tissue | E-MTAB-506, E-MTAB-5060 |

**Figure S1.**
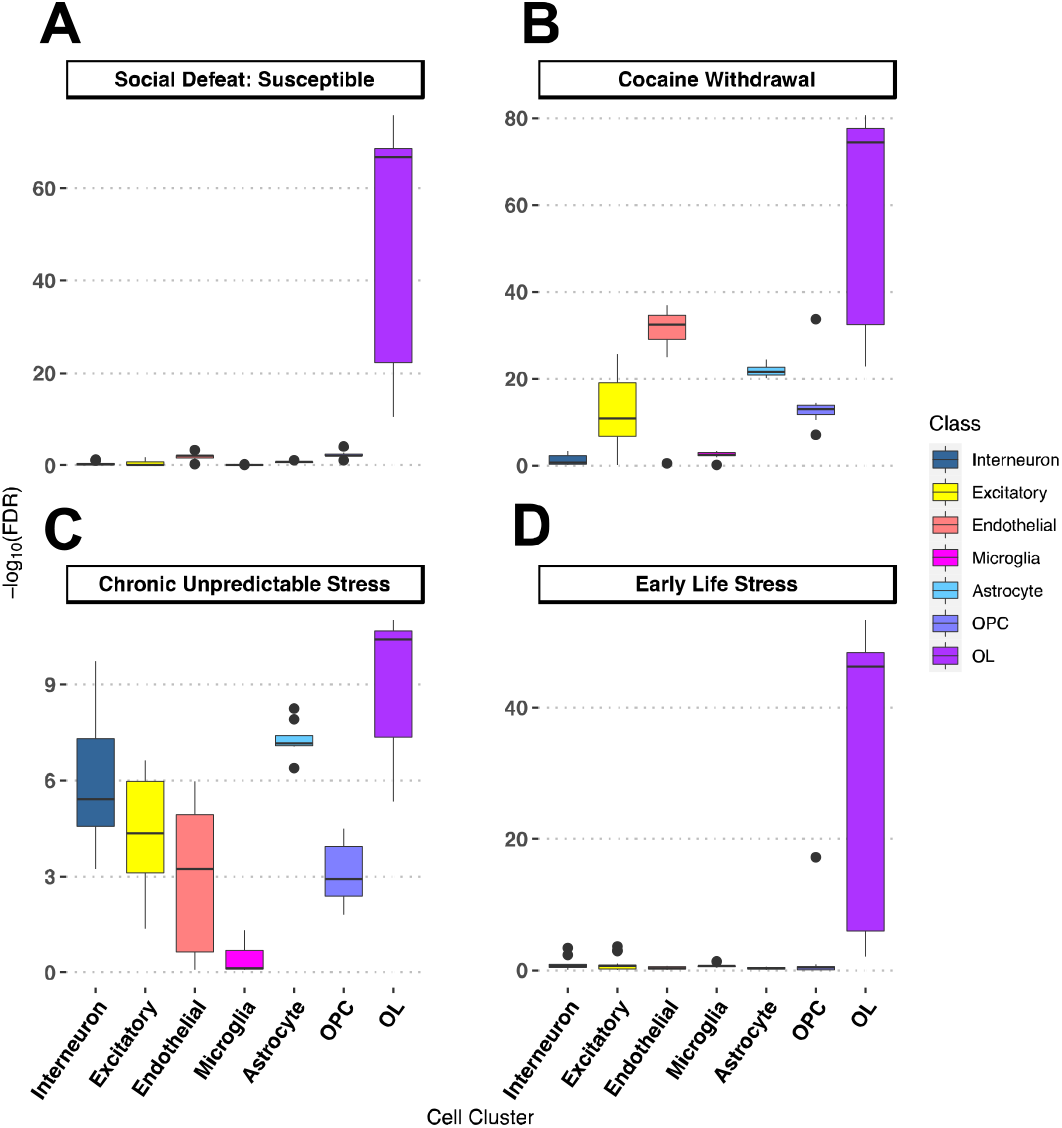
LRcell analyses of animal models of adversity using an additional single cell reference dataset: (A) Susceptibility to social defeat, (B) cocaine withdrawal, (C) chronic unpredictable stress, and (D) early life stress.

**Figure S2.**
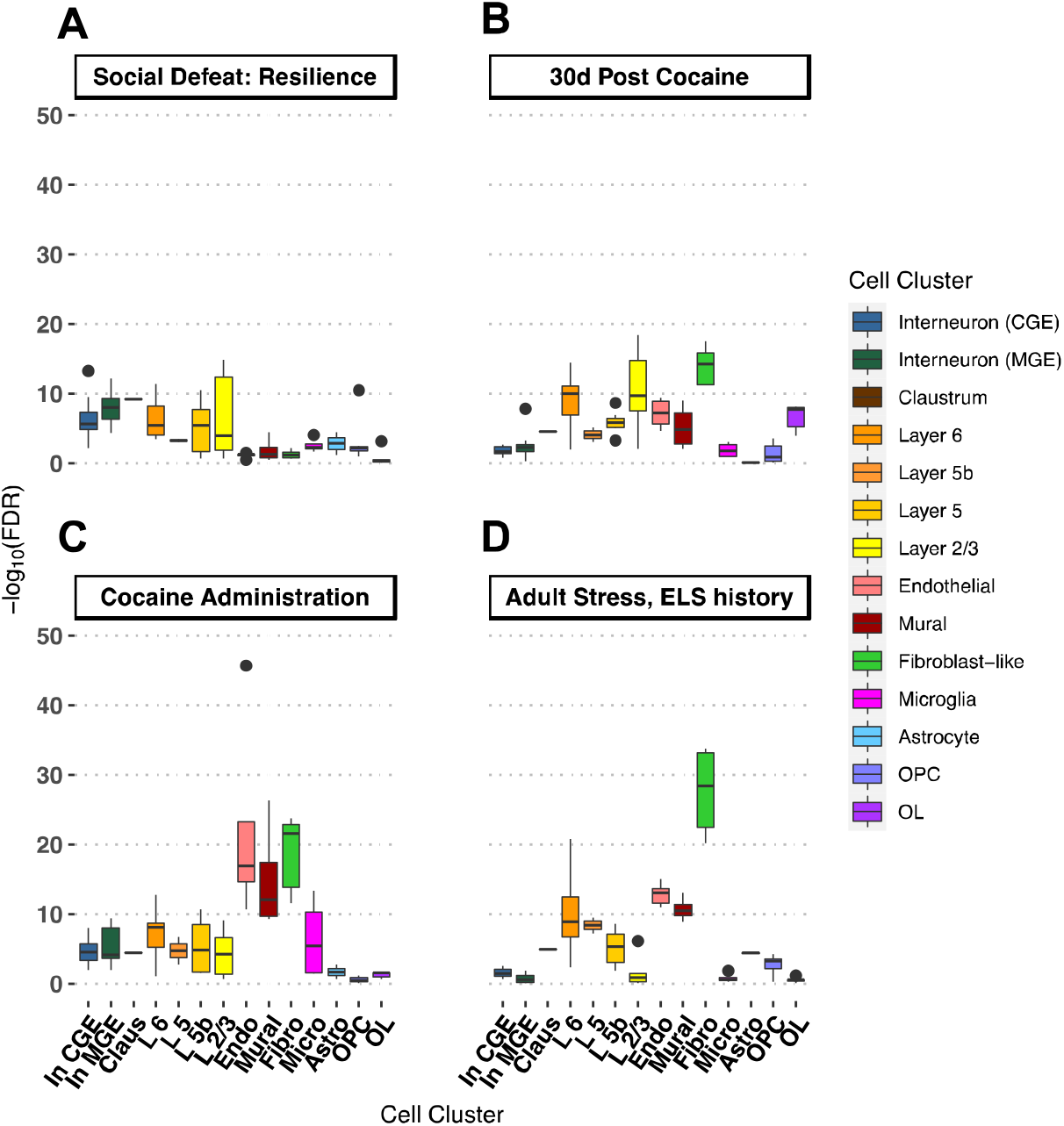
Additional LRcell analyses of animal models. **(A)** Resilience to social defeat stress**. (B)** Thirty days after cocaine self-administration. **(C)** After acute cocaine administration. **(D)** After acute stress in adulthood in animals with a history of early life stress.

**Figure S3.**
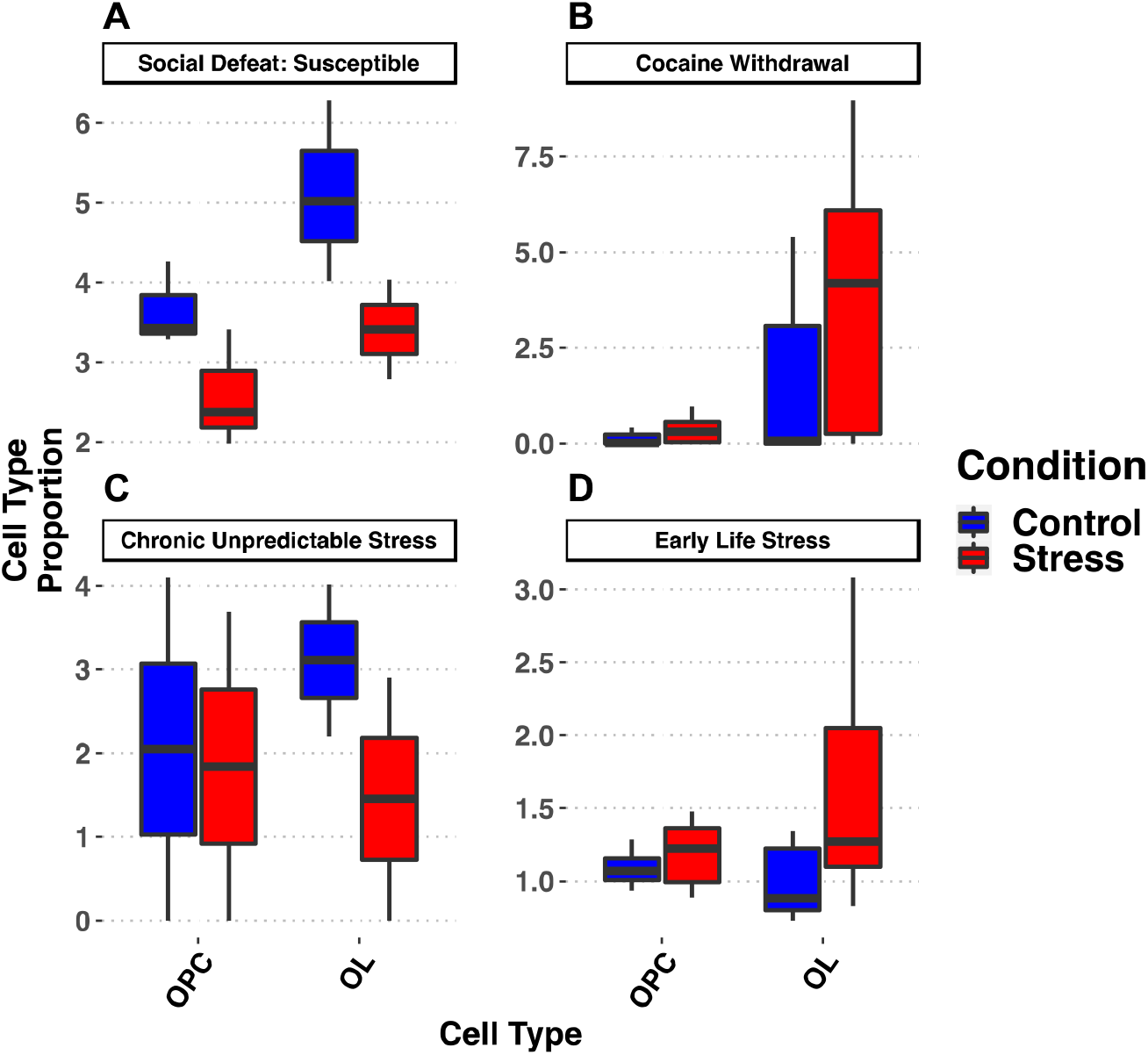
MuSiC deconvolution estimates of OPC and OL proportions across adversity paradigms. Statistical testing was performed using the Welch two-sample t-test for each comparison: **(A)** susceptibility to social defeat (OPC: t(3.6) = 2.05, p-value = 0.12, n= 3; OL: t(3.1) = 2.23, p-value = 0.10, n = 3); **(B)** cocaine withdrawal (OPC: t(92.41) = −4.92, p-value = 4.33 × 10^-6^,n = 56; OL: t(92.4) = −4.77, p-value = 6.93 × 10^-6^, n = 56); **(C)** chronic unpredictable stress (OPC: t(1.98) = 0.07, p-value = 0.95, n = 2; OL: t(1.68) = 0.97, p-value = 0.45, n = 2); **(D)** early life stress (OPC: t(7.61) = −0.91, p-value = 0.39, n = 6; OL: t(6.0) = −1.73. p-value = 0.13, n = 6).

**Figure S4.**
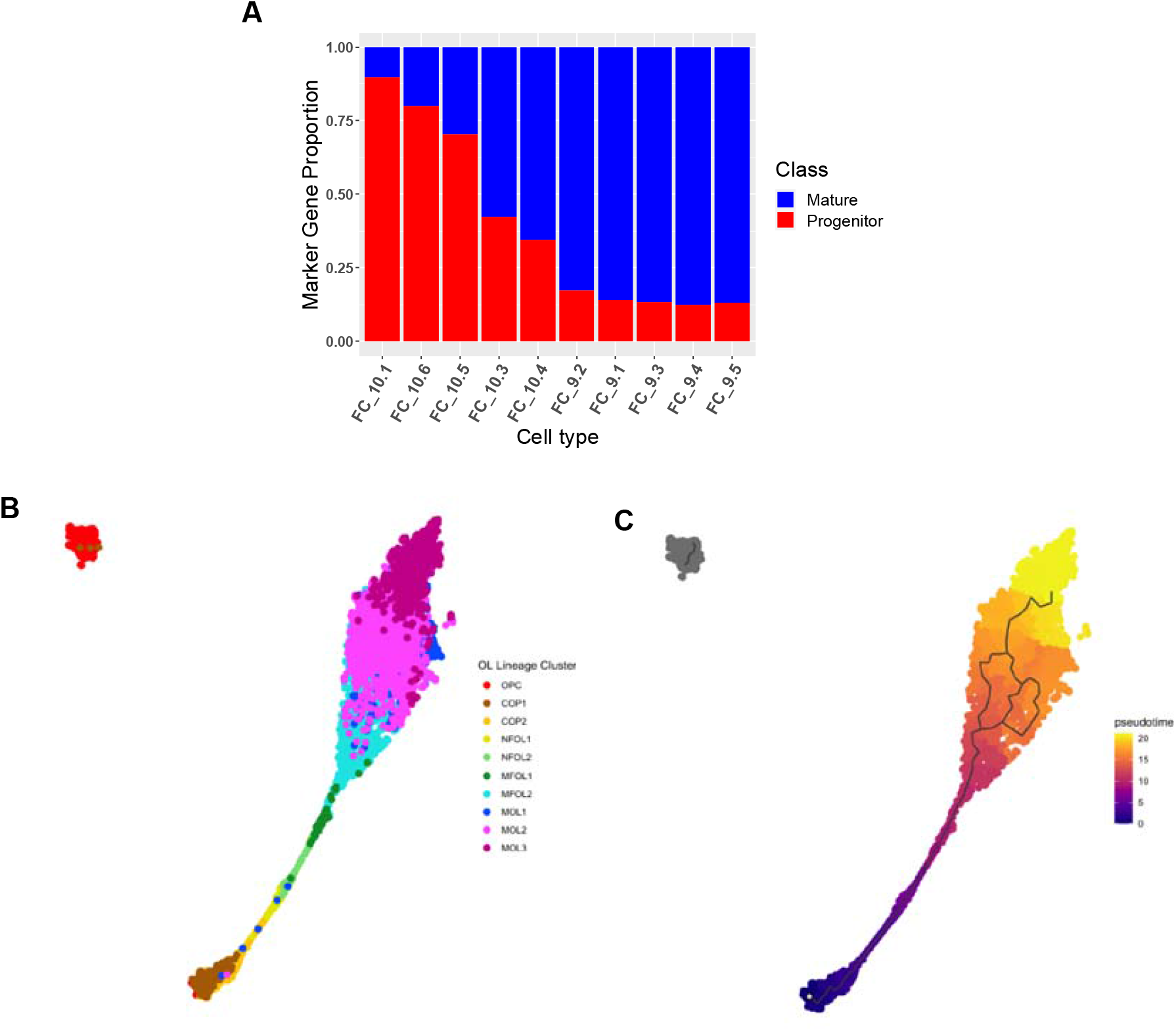
Oligodendrocyte maturation trajectory analysis. **(A)** Proportion of marker genes overlapping with the cell marker database for each OPC and OL cell-type. Bars are colored according to the percentage of overlap between Cell Marker genes classified as mature and progenitor. UMAP trajectory analysis colored by **(B)** cell types according to expression of known marker genes, or **(C)** pseudo-timecourse.

**Figure S5.**
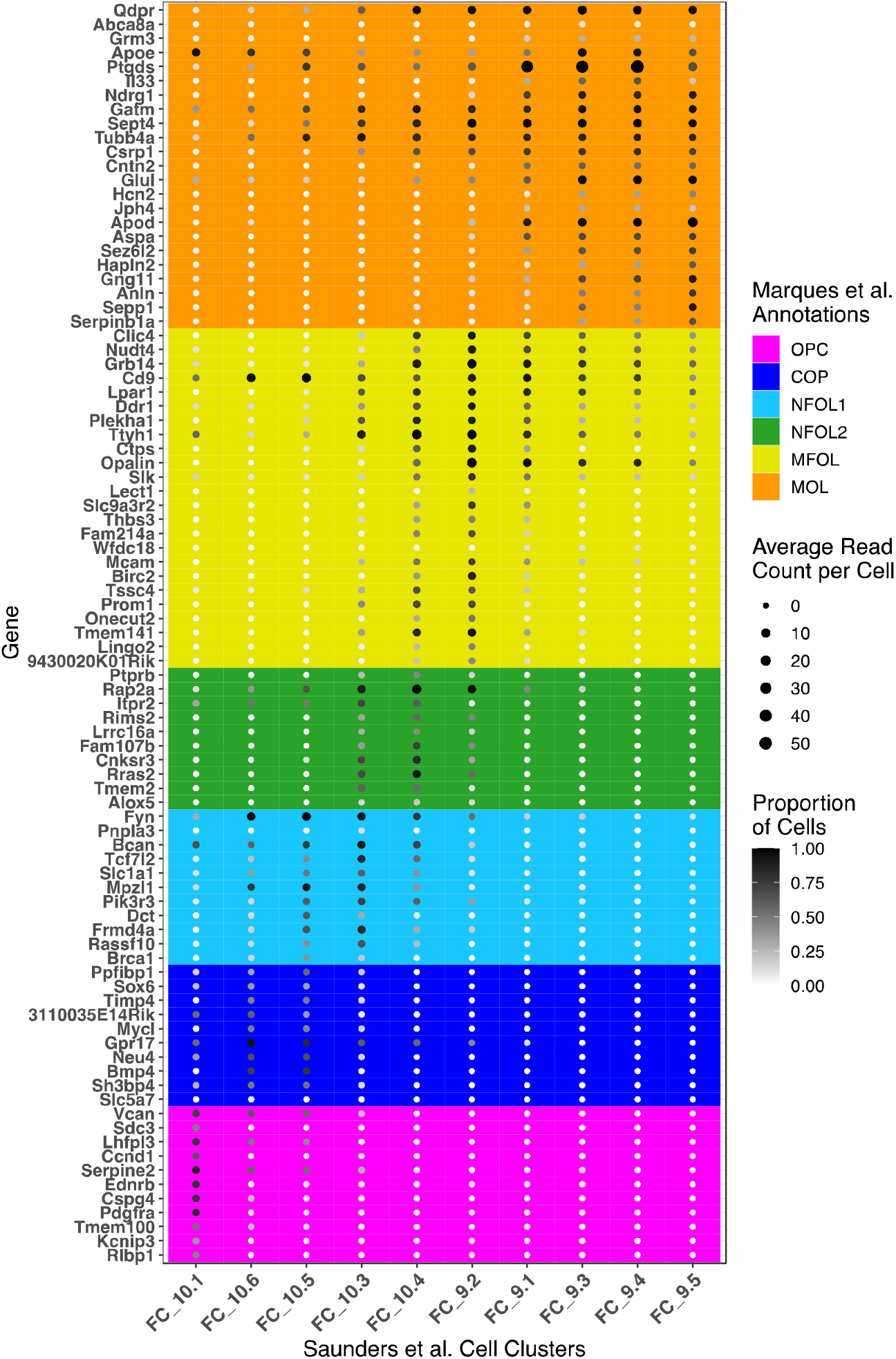
Expression of known maturation stage-specific genes (y-axis) in OL-lineage cell types (x-axis). Marker gene categories are colored according to the maturation stage they define. Dot size represents average gene counts per cell in defined cell types, and dot shade represents the proportion cells expressing the gene. ***Abbreviations:* OPC**: oligodendrocyte precursor cell, **COP**: committed oligodendrocyte precursor, **NFOL**: newly formed oligodendrocyte, **MFOL**: myelin forming oligodendrocyte, and **MOL**: mature oligodendrocyte.

**Figure S6.**
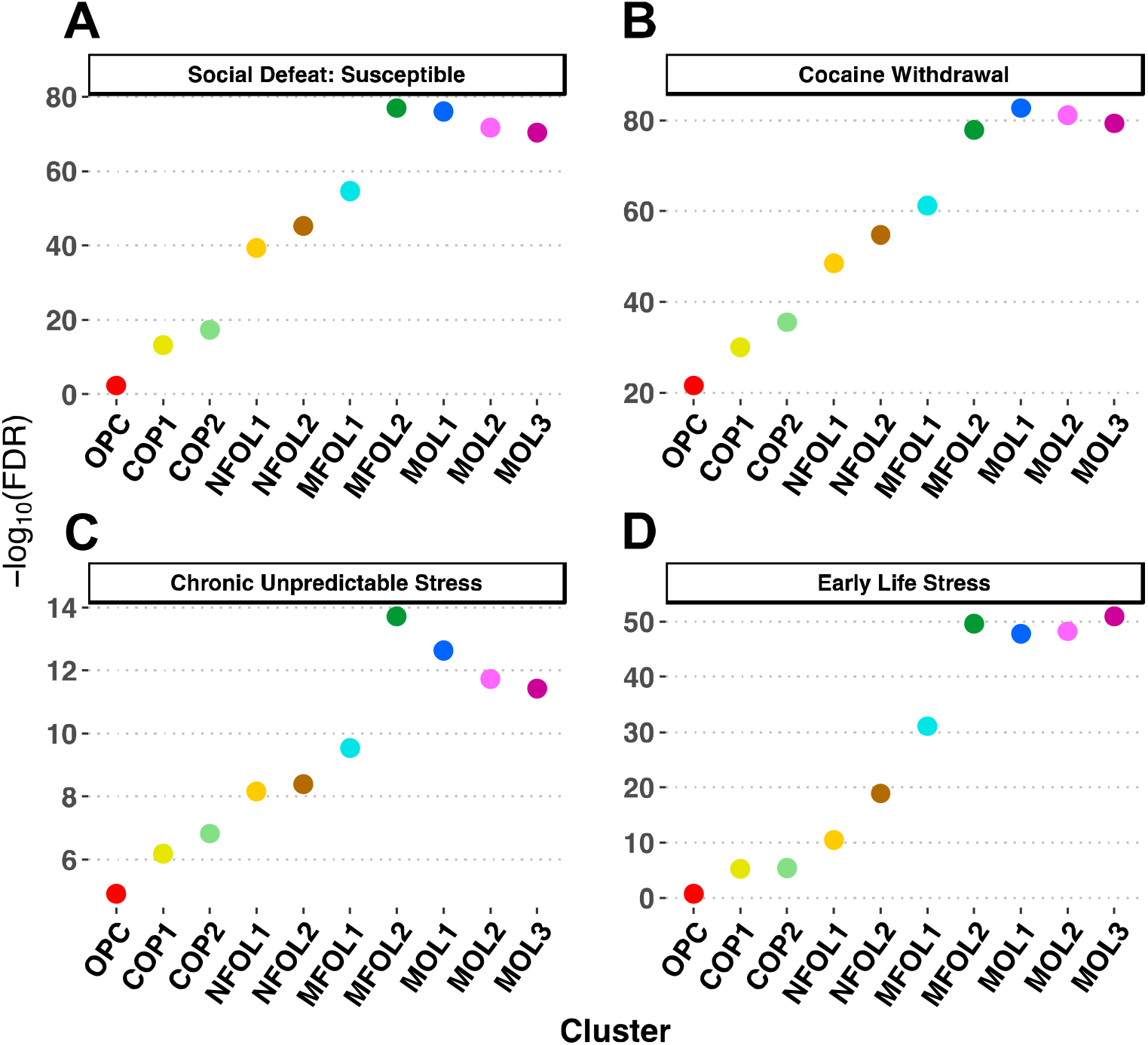
LRcell analysis of individual OL-lineage cell types across adversity paradigms. LRcell analyses of OL cell-types from differential expression analyses of mice (A) susceptible to social defeat, (B) undergoing acute withdrawal from cocaine, (C) after 30 days of chronic unpredictable stress and (D) with a history of early life stress (maternal separation and limited bedding). OL cell-types are ordered from least to most mature. **OPC**: oligodendrocyte precursor cells; **COP**: committed oligodendrocyte precursors; **NFOL**: newly formed oligodendrocytes; **MFOL**: myelin forming oligodendrocytes, **MOL**: mature oligodendrocytes.

**Figure S7.**
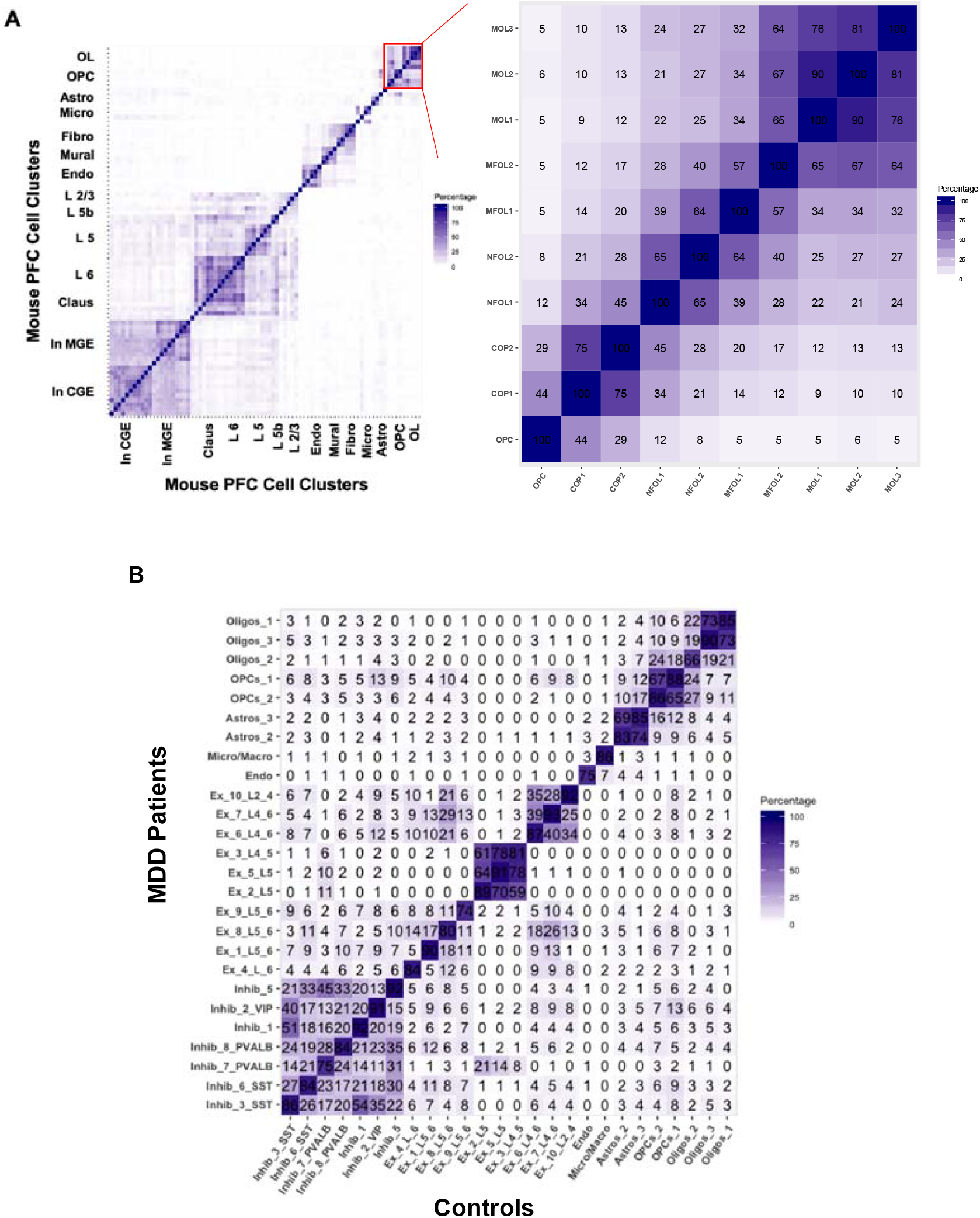
Concordance between marker gene sets. **(A) Left:** Self-correlation of cell-type marker genes in the mouse PFC and **Right:** expansion of OPC and OL populations. **(B)** Overlap in marker genes calculated from human MDD patients (y-axis) or healthy controls (x-axis).

